# CrosstalkNet: mining large-scale bipartite co-expression networks to characterize epi-stroma crosstalk

**DOI:** 10.1101/102848

**Authors:** Venkata Manem, George Adam, Tina Gruosso, Mathieu Gigoux, Nicholas Bertos, Morag Park, Benjamin Haibe-Kains

**Author notes:** Co-first authors.

## Abstract

**Background:** Over the last several years, we have witnessed the metamorphosis of network biology from being a mere representation of molecular interactions to models enabling inference of complex biological processes. Networks provide promising tools to elucidate intercellular interactions that contribute to the functioning of key biological pathways in a cell. However, the exploration of these large-scale networks remains a challenge due to their high-dimensionality.

**Results:** CrosstalkNet is a user friendly, web-based network visualization tool to retrieve and mine interactions in large-scale bipartite co-expression networks. In this study, we discuss the use of gene co-expression networks to explore the rewiring of interactions between tumor epithelial and stromal cells. We show how CrosstalkNet can be used to efficiently visualize, mine, and interpret large co-expression networks representing the crosstalk occurring between the tumour and its microenvironment.

**Conclusion:** CrosstalkNet serves as a tool to assist biologists and clinicians in exploring complex, large interaction graphs to obtain insights into the biological processes that govern the tumor epithelial-stromal crosstalk. A comprehensive tutorial along with case studies are provided with the application.

**Availability:** The web-based application is available at the following location: http://epistroma.pmgenomics.ca/app/. The code is open-source and freely available from http://github.com/bhklab/EpiStroma-webapp.

## BACKGROUND

Advances in high-throughput technologies and the continuous inflow of transcriptomic data have created new avenues to understand complex processes in biological systems. These data repertoires have been used to build networks that aims at decoding complex traits of genetic events through data-driven hypotheses. Biological networks are considered as a promising way to decrypt the molecular interactions between various phenotypes. In this regard, network visualization is a powerful tool to mine, interpret and analyze large sets of biological interactions. A premise that has gained importance in the last few years in designing novel anticancer therapeutics is to target the tumor microenvironment. Tumors interact with the microenvironment and recruit surrounding stromal cells, and depend on these cells for progression, invasion, resistance to therapies; thus this interaction constitutes a major contributor to the carcinogenesis [1,3]. During the last decade, we came to recognize that one of the most important signaling networks in tumor biology is the communication between tumor epithelial (epi) cells and its neighboring stromal cells [1–3]. The interaction between the tumor epi and stromal cells is regulated by a dynamic network of growth factors, cytokines and chemokines. The complexity of these inter and intra-cellular communications is mediated through regulatory loops between the tumor epithelial and stromal compartments. Despite the importance of epithelial-stromal crosstalk in the evolutionary dynamics of cancer, little is known regarding the key players that orchestrate tumor growth, progression, invasion and several other biological processes. A promising approach to shed light on these key players lies in constructing tumor epi-stromal co-expression networks using the transcriptional profiles of epithelial and stromal sample pairs [4–8]. Given the computational and biological complexity of the co-expression network and the number of possible epi-stroma interactions, however, it is challenging to explore and comprehend to these large-scale networks.

Epi-stromal co-expression networks can be represented as a bipartite network where each node represents a gene and each edge denotes a co-expression relationship between the epithelium (epi) and stroma (Figure 1). Throughout this manuscript, we use edge weight, or interaction strength to refer to a co-expression relationship characterized by the underlying correlation between the expression of genes in the epithelial and stromal compartments. For the sake of clarity, relationships between genes within a compartment are not depicted in the epi-stroma interaction network. From a biological point of view, a self-loop is defined as those genes that are co-expressed in both the epithelium and stroma, and from a network point of view, self-loops are all the diagonal elements in an adjacency matrix (defined as a matrix, **A**, where each element **A**[i,j] represents an edge between nodes i and j of a given graph; Figure 1A). These self-loops could potentially be due to an external stimulus in the tumor-stroma macroenvironment, such as hypoxia, or, due to paracrine interactions caused by several growth factors. For instance, in Figure 1A, the G2 gene in epithelium is connected to the G2 gene in the stroma, therefore forming a self-loop. Knowing the neighborhood of gene(s) is also critical to any biological network. For instance, the first neighbors of the G4 gene in epi are G1, G5 in the stroma, while the second neighbors of the G4 gene in epi are all the connections of G1, G5 in the stroma, which is G5 in epi. The biological function of a node is determined by all its interactors, or neighbors. Some genes are highly connected (hubs) and could be regarded as the most important nodes or influencers in the network. For instance, the master regulators in regulatory networks correspond to the highly connected nodes in the network [9].

**Figure 1:**
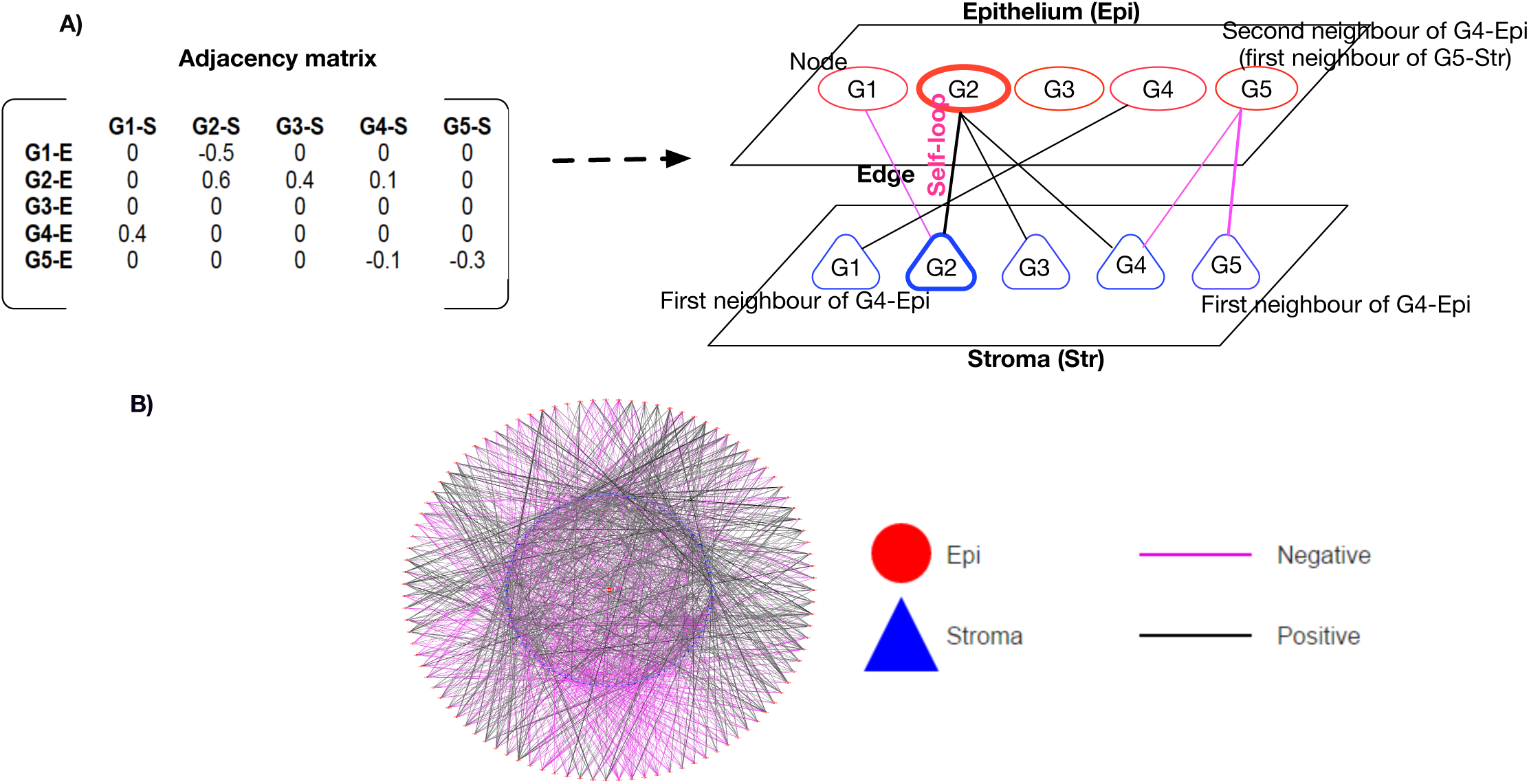
Co-expression of genes in tumor epithelial and stromal cells. **(A)**Schematic diagram of a co-expression network (which is a bipartite graph) between tumor epithelium and stroma components with 5 genes along with the adjacency matrix. Panel B-Genome wide epi-stroma (sparse) interaction network. (**B**)Red, Blue nodes represent epi and stroma genes respectively. The color gradient of the edges denote the strength of the co-expression relationship. Black, magenta colors denote positive and negative co-expression relationships between the genes, respectively.

Differential networks [10] are used to characterize the rewiring of epi-stroma interactions in the tumor specific network, as shown in Figure 2. This network uncovers the interactions gained and lost by the tumor during its evolution from a normal state.

**Figure 2:**
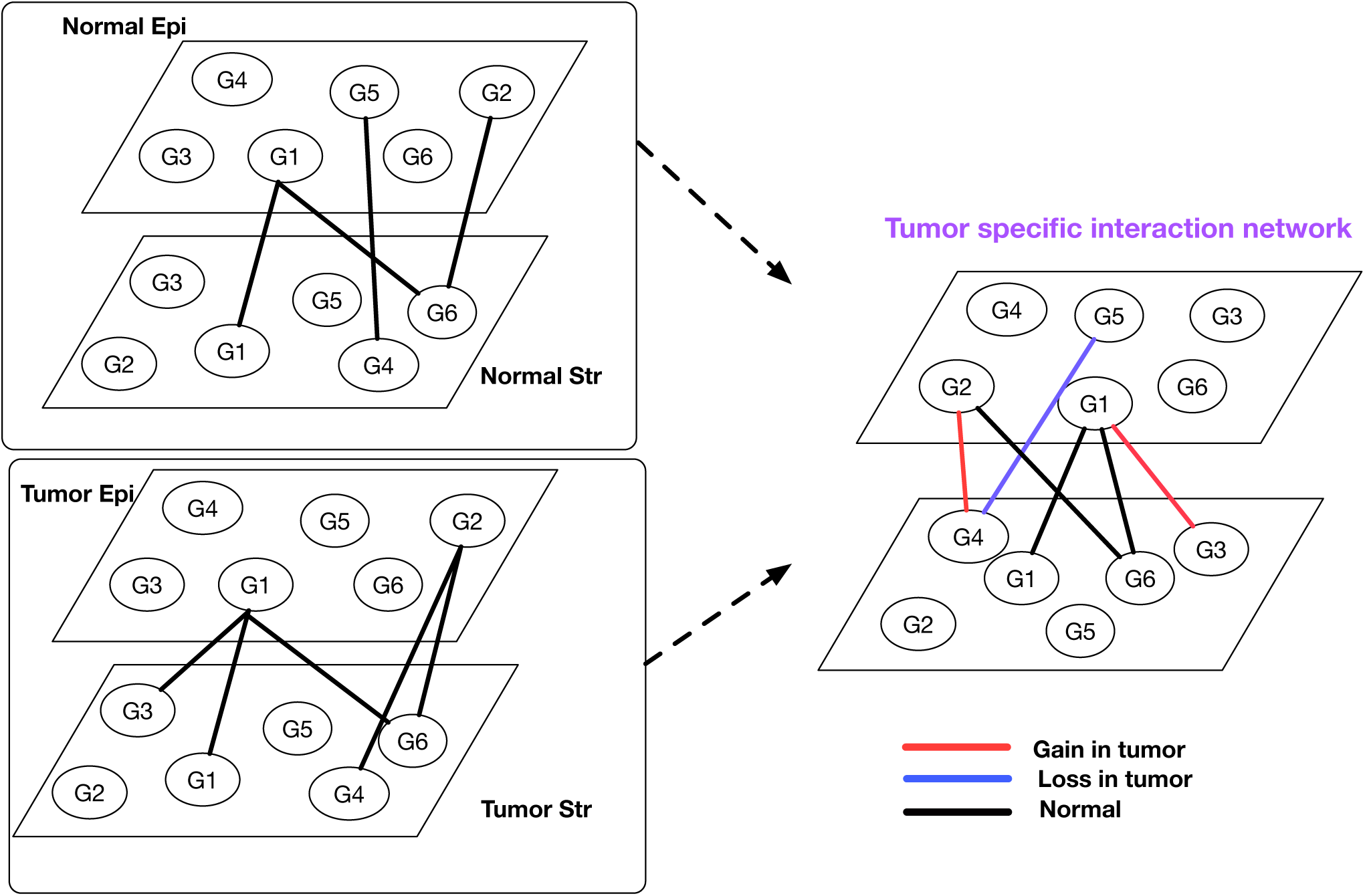
Design for identifying the tumor specific epi-stroma interactions, using the notion of a differential network. A schematic diagram of a differential network inferred from matched normal and tumor epithelium and stromal pairs.

A variety of tools have been developed in the literature to mine large-scale interaction graphs in the context of biological networks [11–19]. Due to the advancement of web-based technologies, web-applications have been widely used to facilitate the mining and rendering of large scale graphs interactively [20–22]. Although these web-based tools enable retrieval of putative gene–gene interactions, current web-based applications do not support the visualization of differential co-expression bipartite networks. This is particularly important in biological interactions such as the crosstalk between tumor epithelial and stromal cells. To circumvent the above limitation, we developed *CrosstalkNet*, a web-based tool for exploratory analysis of large-scale bipartite networks. We believe that this web-based application will assist researchers across disciplines in mining co-expression networks to unravel novel biological insights into the epithelial-stroma crosstalk that occurs in tumors. In addition, this approach is applicable to other systems where interactions between two compartments are of biological importance.

### Implementation

Systematically mining and visualizing large-scale graphs for meaningful biological inferences remains one of the main challenges in network biology. To address this issue we developed *CrosstalkNet*, a web-application enabling users to upload their own inferred bipartite networks and mine these large graphs through their web browser. To ensure rapid rendering of graphs, *CrosstalkNet* limits the visualised networks to 10,000 elements (comprising of both nodes and edges) per graph. At any time, users can export the figures in a high-resolution PNG format and tables in a CSV format for data sharing and further manipulations. *CrosstalkNet* is divided into four major panels, namely, *Main graph, Interaction explorer, Path explorer, Degree explorer* (see Figure 3). The functionalities of each panel are described below in detail. Case studies will be further presented to exemplify the use of each of the panels.

**Figure 3:**
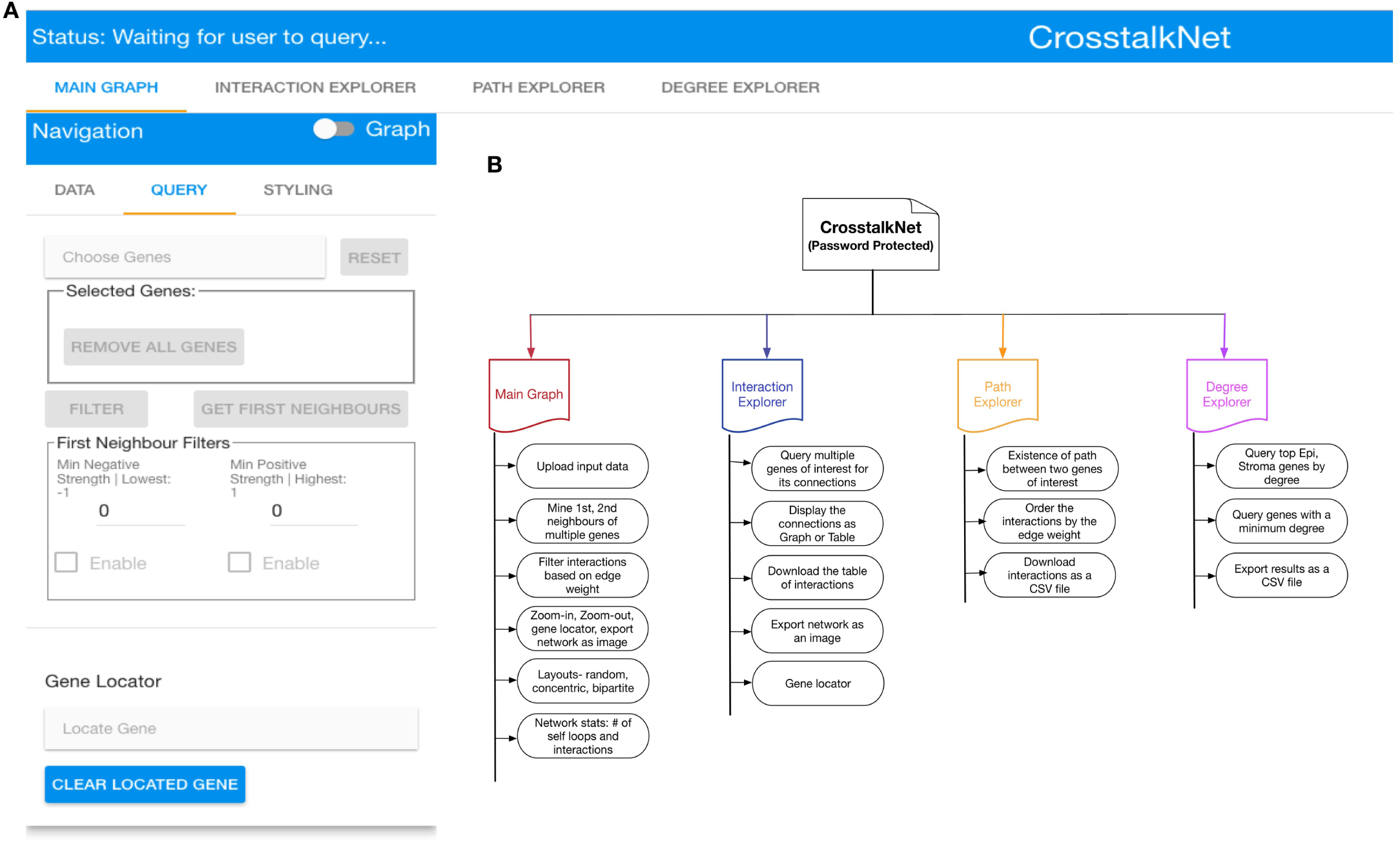
The design of the CrosstalkNet web-application. *CrosstalkNet* allows for guest or password-protected user access. (**A**)Main page of *CrosstalkNet* where users can load their files and start exploring their bipartite networks. (**B**)Flowchart representing each analysis panels. The panel *Main Graph* is used to explore the neighborhood of multiple genes. The panel *Interaction Explorer* is used to explore neighbors for a given gene with infinite hops. The panel *Path Explorer* is used to explore all the existing paths between two genes of interest. The panel *Degree Explorer* is used to explore the connectivity of genes in epi and stroma compartments.

*Main Graph:* One of the important topological features of a network is the neighborhood of interest. Given that the number of interacting genes (network neighbors) grows quickly with the number of “hops”, *CrosstalkNet* enables visualization of the first and second levels to ensure efficient exploration of the network topology around genes of interest. The first level neighborhood of a gene is defined as the set of all genes connected to it, while the second level neighborhood is defined as the set of genes connected to the first level neighborhood genes. Mining the neighbors in a very dense graph is a non-trivial task. Hence, we implemented a filtering function to help users identify the most relevant interactions based on the edge weights, i.e., co-expression of genes between the epi and stroma compartments.

*Interaction Explorer:* Many genes are simultaneously involved in multiple biological pathways, which govern the functional processes of a cell. In this regard, the cells exchange messages through autocrine, paracrine and endocrine signaling pathways, which are the key components in the epi-stroma crosstalk. Autocrine signaling is defined as a mechanism in which the cells send signals to itself, by releasing a ligand that binds to the receptors on its own surface. Paracrine signaling is a mechanism in which the cells send signals to nearby cells, and induce changes to them. Endocrine signaling is a mechanism in which the cells communicate to distant cells through several growth factors. In order to capture these signals, it is important to explore these regulatory loops in a co-expression network, in a systematic manner. For this purpose, we developed the “Interaction explorer” panel that provides the ability to view interactively many levels of neighbors or connections to multiple genes in a bipartite graph.

*Path Explorer:* In a dense (complete) network it is non-trivial to find interactions along with the strength of the co-expression relationship between any two genes of interest. For this purpose, we developed the “path explorer” panel that provides the user with the ability to obtain all paths between two genes of interest with a maximum of one hop.

*Degree Explorer:* One of the important features of a network is the hubs. Hubs are defined as those genes that have highest connectivity compared to other nodes in the network. Since the tumor epi-stroma interaction network is a bipartite graph, the hubs could potentially be different in the epi and stroma compartments. It is important to study the hubs in both the epithelial and stromal compartments, as the network biology and structure depends on the connectivity of nodes in epi as well as in the stroma compartments. We therefore developed the “degree explorer” panel which displays the genes (expressed in epi and stroma) based by their connectivity.

The current implementation of *CrosstalkNet* allows the users to visualize co-expression networks with up to 10,000 elements (including both nodes and edges).

## RESULTS

In this study we show how the *CrosstalkNet* web-application can be used to uncover the epi-stroma co-dependencies in estrogen receptor-positive (ER+) breast cancer patients. We show hereafter how *CrosstalkNet* enables efficient exploration of complex gene interactions through five case studies describing how to (*i*) extract a shared neighborhood (first neighbors) of genes of interest; (*ii*) explore paracrine interactions, (*iii*) mine a differential network, (*iv*) extract paths between any two functionally similar genes; (*v*) extract and mine the hubs in the network.

### Inference of the epi-stroma interaction network

Using published transcriptomic profiles of laser-microdissected samples from our previous study [4] (Supplementary Table 1), we inferred a genome-wide network by estimating pairwise co-expression interactions between the tumor epithelium and tumor stroma along with normal epithelium and normal stroma (see Methods section). These tumor and normal bipartite networks were statistically compared to identify epi-stroma interactions that are specifically gained or lost in the tumors; this set of epi-stroma interactions is referred to as a *differential* network. These networks are intrinsically large-scale as they constitute a graph-based representation of the complex crosstalk that occurs between epithelial and stromal cells in tumor and healthy tissues. In this study, a false discovery rate (FDR) of 15% is applied to select all the significant interactions from the epi-stroma co-expression network. Concordant with our previous study [4], we found that the proportion of self-loops are greater in number in the tumor network than in the normal network (Supplementary Figure 1); a subset of these self-loops was previously validated by analysis of immunohistochemistry [4]. We hypothesize that these self-loops are housekeeping genes specific to the tumor epi-stroma microenvironment, which are expressed in certain pathophysiological conditions. This suggests that functional rewiring of such interactions is an important feature in breast tumorigenesis. Moreover, we observed that the degree distribution of the epi and stroma compartments in the tumor network (Supplementary Figure 2) may follow a power law distribution (with the rejection of null hypothesis: *p*_*epi*_ of 0.57 and *p*_*str*_ of 0.45 respectively). This implies that the network is a scale-free graph and very few genes in the graph have high degree [23]. The advantage of scale-free networks often observed in biological settings [23] is that they are more robust, and random loss of an individual non-hub node is less disruptive to the network compared to a random network. This conveys that a relatively small number of nodes have an important role to play in the epi-stroma interaction network, and thus these could represent potential actionable targets within the tumor epi-stroma network.

### Identification of shared biological property between two genes

Most of the ER+ breast cancer patients treated with aromatase inhibitors develop resistance.. Recent evidence indicates that drug resistance is due to an aberrant function of the interferon signaling pathway, resulting in the amplification of interferon-induced genes [24]. Many studies showed that these genes, which include OAS1 and IFIT1, play an important role in treatment-induced therapy resistance [25–27]. Using the *Neighbor Explorer*, we therefore sought to explore the shared epi-stroma interactions of IFIT and OAS1 in tumor epithelial cells (Figure 4). The similarity of any IFIT1-Epi and OAS1-Epi nodes in a network can be quantified by the number of neighbors that are shared between these two nodes. We found that IFIT1 and OAS1 in epi have 17 and 16 interacting genes in stroma, respectively, out of which 15 genes are common between the two nodes (hypergeometric test p-value < 1E-16). Pathway enrichment analysis was performed on these 15 genes to obtain an insight into the biological processes using the Gene Ontology (GO), KEGG and REACTOME pathway lists. A total of 7 pathways are over-represented with an FDR<5%, which include immune- and interferon-related pathways that regulate the interferon-induced genes. This finding provides additional evidence supporting the importance of the interferon signaling pathway-induced genes in ER+ breast tumors.

**Figure 4.**
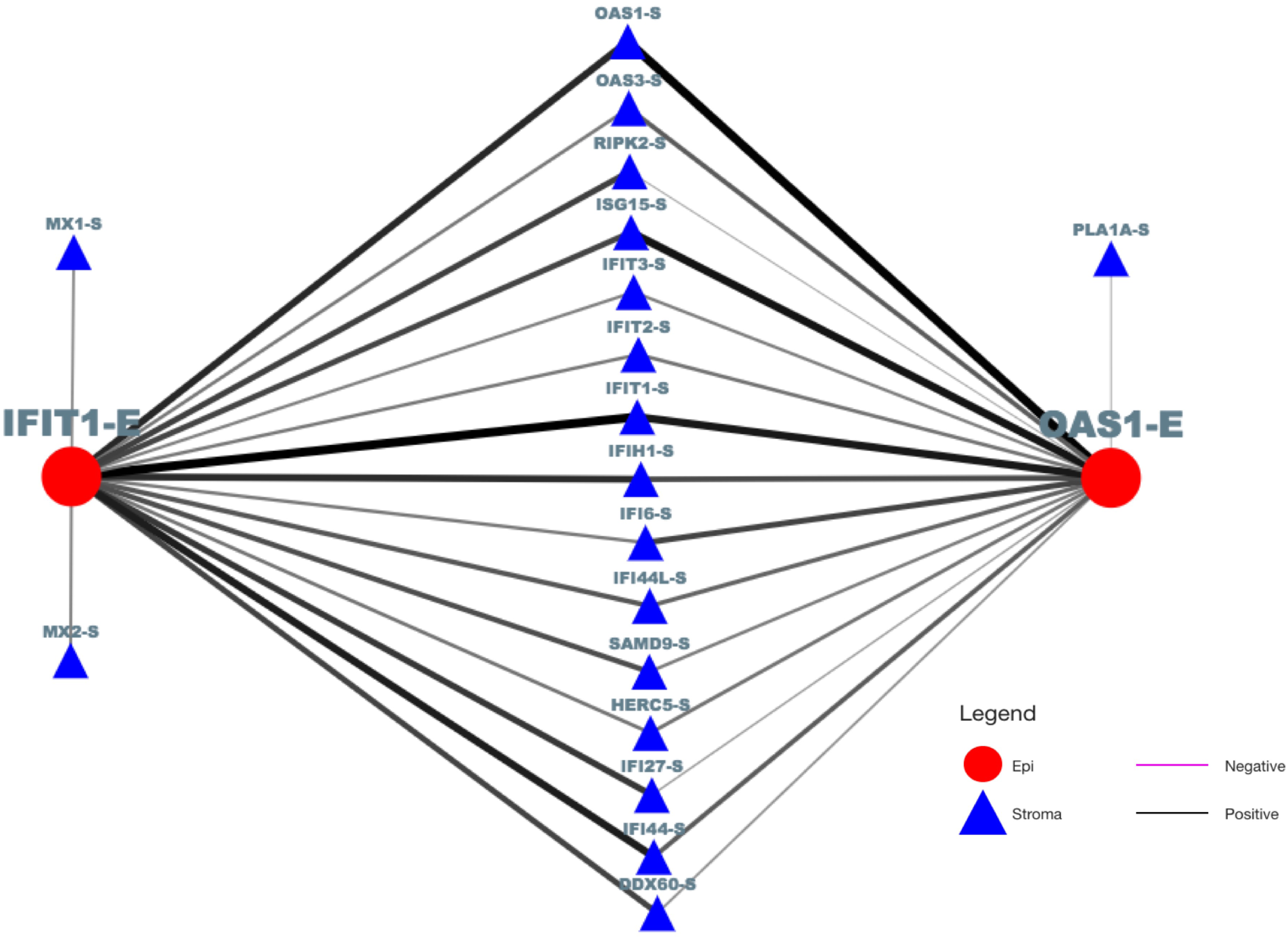
A case study for the *Neighbor Explorer* panel. Shared Neighborhood between theIFIT1 and OAS1 genes in tumor epithelial cells. “-E” and “-S” refer to gene expression in epithelial and stromal cells, respectively. Thickness of the edge indicates the strength of interaction, i.e., correlation. Black and magenta edge colors denote positive and negative correlation, respectively.

### Identification of paths between functionally similar genes

A path is defined as a co-expression relationship between an epithelial gene and a stromal gene. Therefore, a path between two different epithelial genes involves a path from the first epithelial gene to a stromal gene, and an additional path from that stromal gene to the second epithelial gene. It is well known that proteins interact, modify and co-regulate the gene expression interactions under several micro- and macro-environmental conditions [28]. In this context, functionally-related genes play an important role in characterizing the interactions in the network. For this purpose, users have the ability to mine all the existing paths between any two functionally similar nodes in the network using the path explorer. For instance, the interferon-induced proteins IFIT1 and OAS1 in the tumor epithelium compartment have 15 paths between them through the stromal genes. This can be seen in Table 1, where each row of the table represents a path between IFIT1-Epi and OAS1-Epi. The path explorer panel displays the paths and the corresponding strength of relationship between these functionally similar genes, which are over-represented in interferon and immune signaling pathways. The strengths of the co-expression relationship can provide insight as to whether the intermediate genes are strongly or weakly correlated along with the direction of relationship with the IFIT1 and OAS1 genes.

**Table 1:**
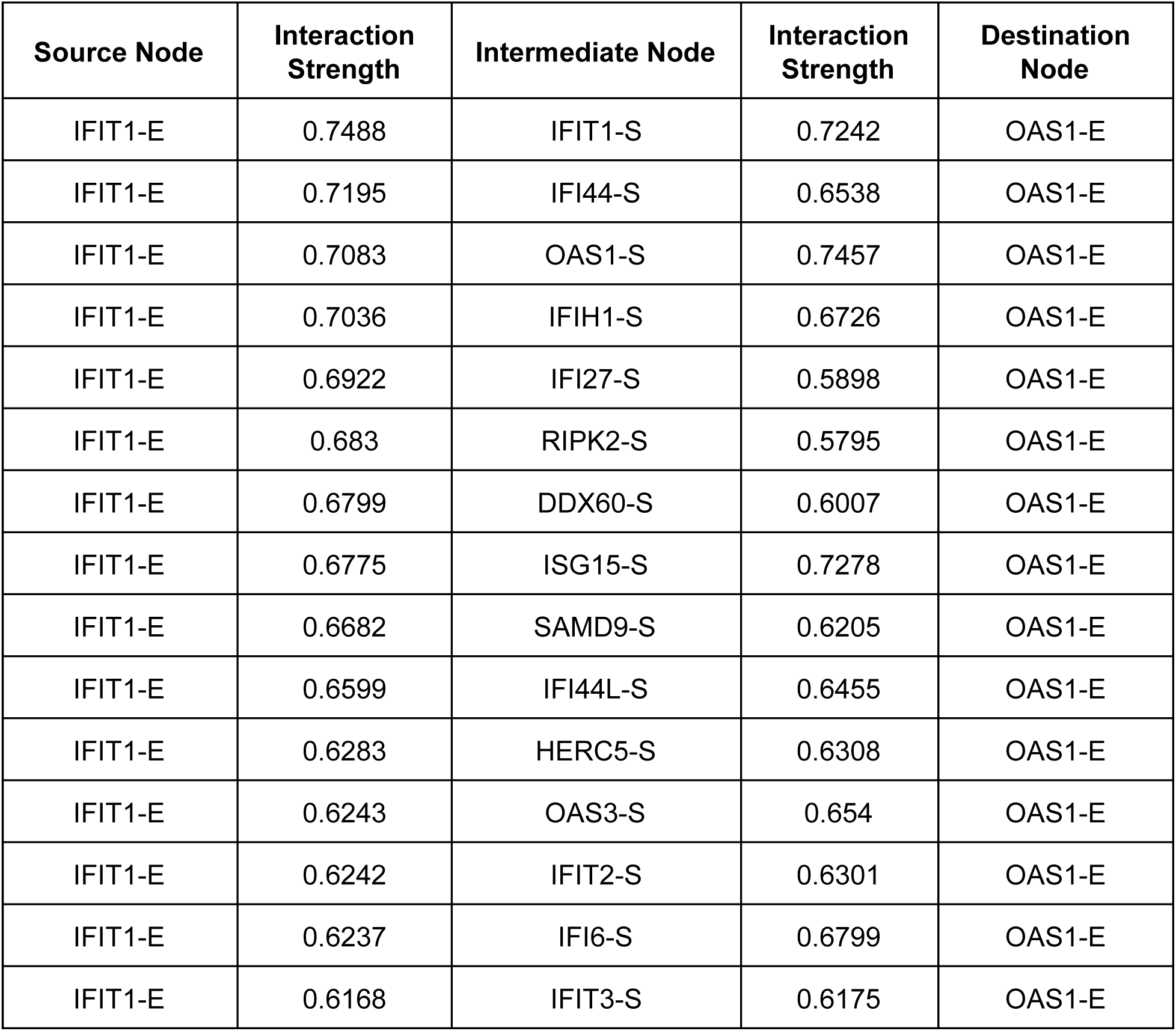
All paths between IFIT1-E and OAS1-E ordered by the interaction strength between IFIT1-E and its corresponding stromal neighbors. “-E” and “-S” refer to gene expression in epithelial and stromal cells, respectively.

| Source Node | Interaction Strength | Intermediate Node | Interaction Strength | Destination Node |
| --- | --- | --- | --- | --- |
| IFIT1-E | 0.7488 | IFIT1-S | 0.7242 | OAS1-E |
| IFIT1-E | 0.7195 | IFI44-S | 0.6538 | OAS1-E |
| IFIT1-E | 0.7083 | OAS1-S | 0.7457 | OAS1-E |
| IFIT1-E | 0.7036 | IFIH1-S | 0.6726 | OAS1-E |
| IFIT1-E | 0.6922 | IFI27-S | 0.5898 | OAS1-E |
| IFIT1-E | 0.683 | RIPK2-S | 0.5795 | OAS1-E |
| IFIT1-E | 0.6799 | DDX60-S | 0.6007 | OAS1-E |
| IFIT1-E | 0.6775 | ISG15-S | 0.7278 | OAS1-E |
| IFIT1-E | 0.6682 | SAMD9-S | 0.6205 | OAS1-E |
| IFIT1-E | 0.6599 | IFI44L-S | 0.6455 | OAS1-E |
| IFIT1-E | 0.6283 | HERC5-S | 0.6308 | OAS1-E |
| IFIT1-E | 0.6243 | OAS3-S | 0.654 | OAS1-E |
| IFIT1-E | 0.6242 | IFIT2-S | 0.6301 | OAS1-E |
| IFIT1-E | 0.6237 | IFI6-S | 0.6799 | OAS1-E |
| IFIT1-E | 0.6168 | IFIT3-S | 0.6175 | OAS1-E |

### Identification of paracrine interactions

Many clinical and biological studies have supported the role of diverse paracrine, autocrine, and endocrine signaling interactions in the tumor ecosystem [3]. It is established that spatial and temporal changes in the tumor growth and invasion occur due to the secretion of cytokines and growth factors in the tumor microenvironment. In this regard, the family of fibroblast growth factors (FGFs) plays an important role in proliferation, invasion and metastasis in breast cancer [29,30]. In addition, recent studies have supported FGF as a potential target in breast cancer [31]. Using the *Interaction Explorer*, users have the ability to mine the paracrine or autocrine interactions between the tumor epithelial and stromal cells. Our results suggest that the expression of FGFR1 gene in the stroma sends messages to the epithelium compartment through paracrine signaling, which in turn may trigger FGFR1 gene in the epithelial compartment (Figure 5). This is in agreement with the observation made in [30]. In addition, the *Interaction Explorer* panel allows the user to mine the network to various levels, from which one can obtain neighbors of any order. This functionality enables construction of a path with a specified number of hops from the source gene.

**Figure 5.**
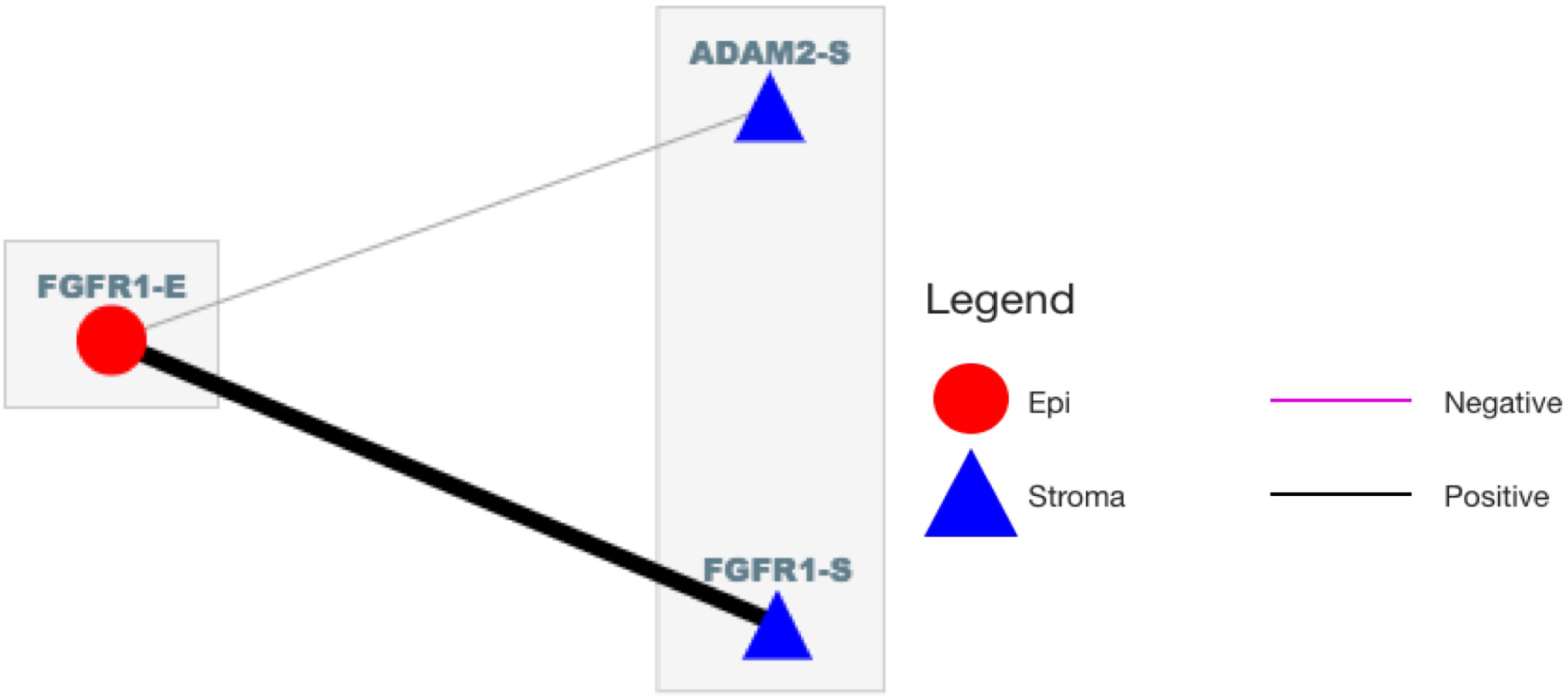
A case study for the *Interaction Explorer*panel. Paracrine interaction of FGFR1-Sand FGFR1-E depicting all the connections of FGFR1-E. “-E” and “-S” refer to gene expression in epithelial and stromal cells, respectively. The thickness of edges indicates the strength of interaction (correlation denoting gene co-expression). Black and magenta edge colors denote positive and negative correlations, respectively.

### Identification of gain/loss of interactions in the tumor epi-stroma network

To identify the epi-stroma interactions that are gained or lost in ER+ breast tumors, we used the *Delta* option in the main graph panel to explore a differential network where each edge strength is defined as the difference of co-expression computed from the tumor and normal epi-stroma interaction networks. The significance of the difference in correlation coefficients was computed using Fisher’s *Z* transformation for two independent samples, followed by FDR correction.

With the tumor-specific epi-stroma (differential) network in hand, we explored the interactions involving S100A7, a protein in the S100 family known to be a key contributor to the onset of aggressive and invasive tumors with the help of the tumor stroma [32]. The changes in the co-expression of S100A7-epi and S100A7-stroma between normal and tumor networks is captured in Figure 6. The strong delta value (△=0.99, p= 6.5E-9) indicates that there is a significant gain of interaction in the tumor. This supports the fact that S100A7 is one of the key genes that is differentially co-expressed between normal epi-stroma and tumor epi-stroma, and contributes to tumorigenesis [32].

**Figure 6.**
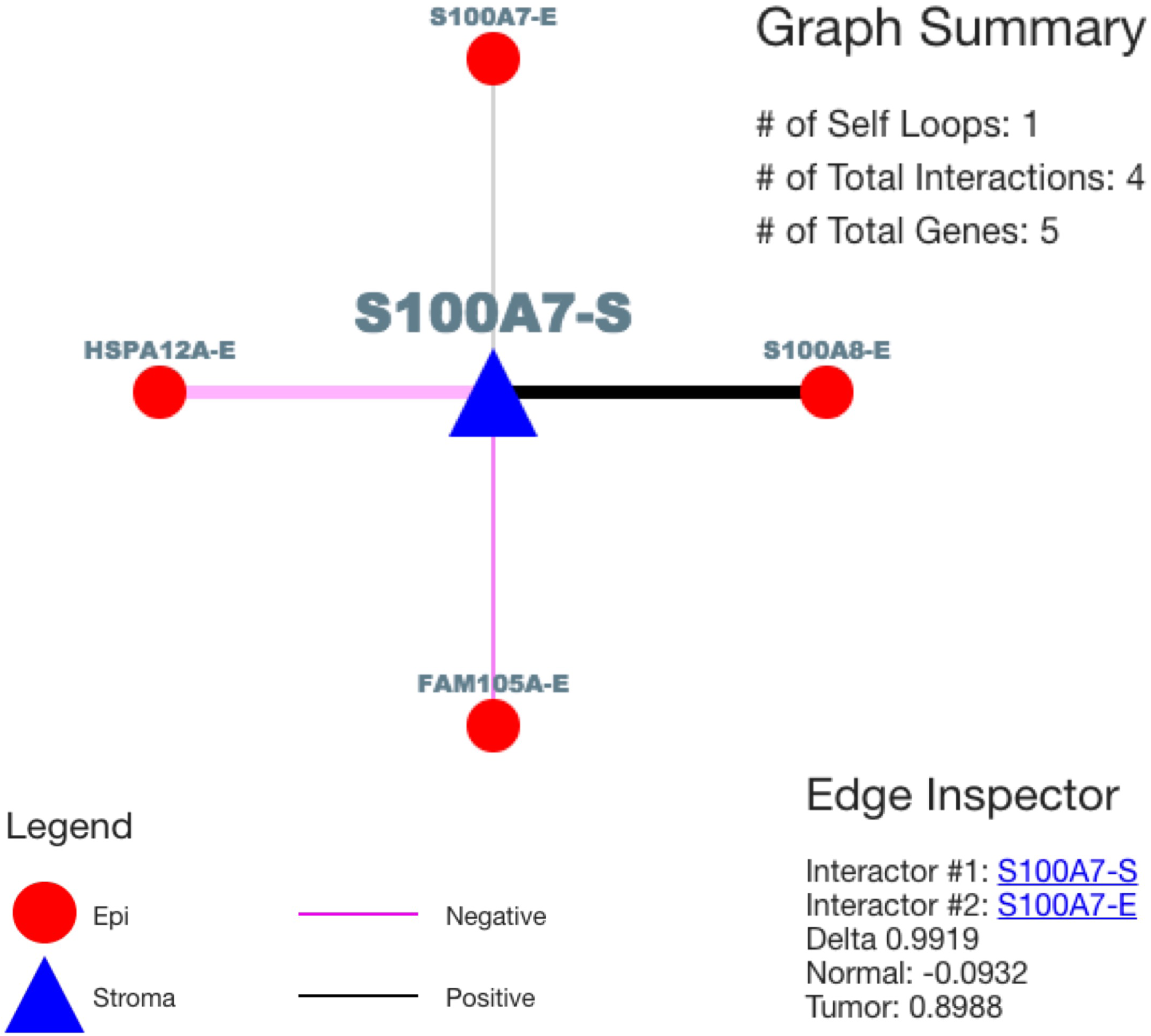
A case study for differential network. Delta network of S100A7 (highlighted in green) genes in ER+ breast tumors. “-E” and “-S” refer to gene expression in epithelial and stromal cells, respectively. The edge inspector gives the difference in the edge weights between tumor and normal, indicating that this interaction is gained in tumor. The thickness of the edge indicates the strength of interaction, i.e. correlation. Black and magenta edge colors denote positive and negative correlations, respectively.

### Identification of hubs (potential drug targets)

The genes with the highest degree in a network, referred to as hubs, are of particular interest, since they involve the most interactions and may therefore play a key biological role in the epi-stroma interaction network. We used the *Degree Explorer* panel to extract the 10 hubs in the tumor epi-stroma network, with genes involved in over 50 interactions (Table 2). The most highly connected gene in the tumor network is the BDNF gene with 71 stromal connections and 3 epithelial connections. Several studies have shown that BDNF enhances tumor progression in breast cancer patients [33,34] and is also a potential target in the treatment of cancer [35].

**Table 2.**
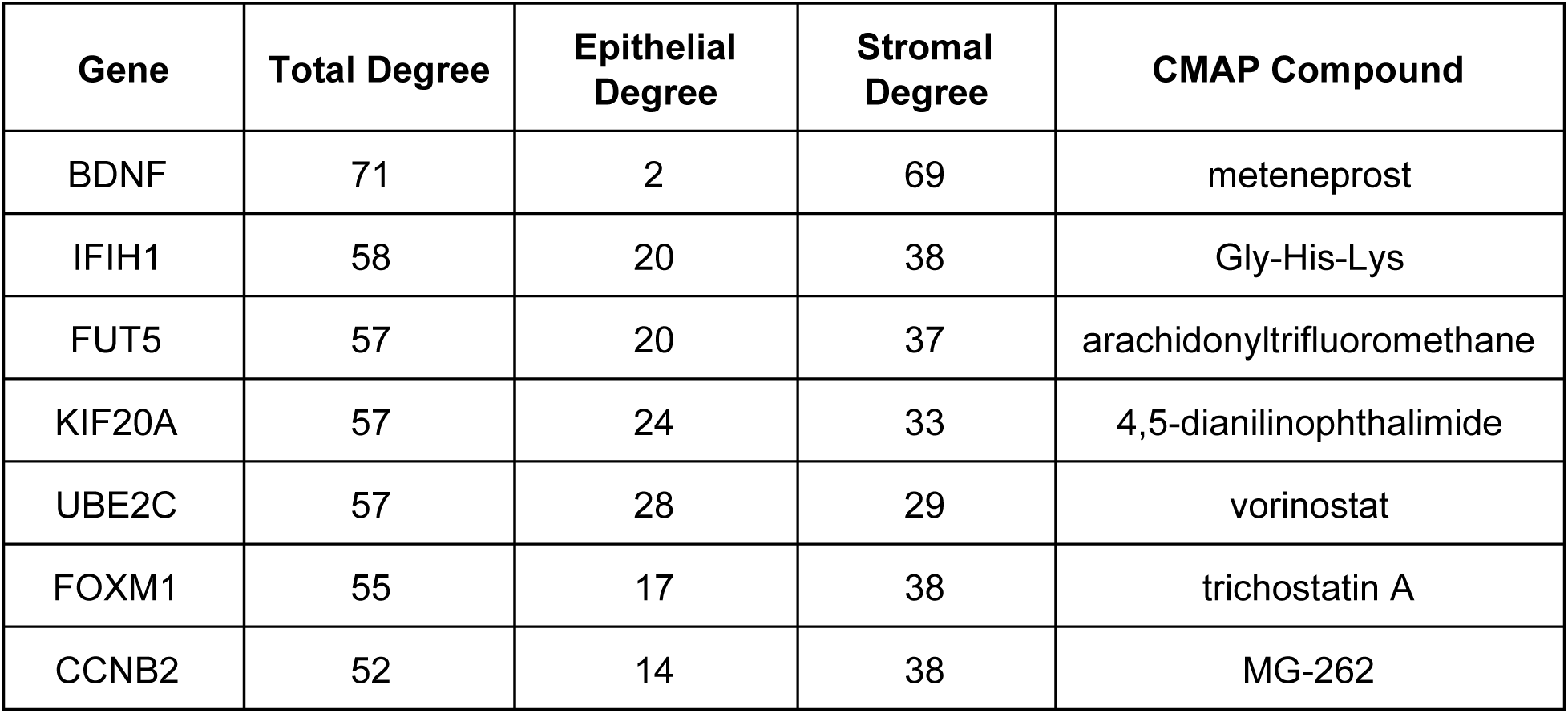
The most highly connected genes in the tumor epi-stroma interaction network involved in more than 50 interactions. Genes are ranked based on their number of first neighbors (epi and stromal connections). The CMAP compound list the drugs that yield the strongest perturbation of the hub gene of interest.

We then leveraged our recent *PharmacoGx* package [36] to characterize the gene expression changes induced by treatment of drugs investigated in the Connectivity Map (CMAP) [37]. We then predicted compounds that specifically target the hubs by inhibiting them in the epi-stroma interaction network (Table 2). Although targeting genes responsible of tumor-specific interactions between epithelial and stromal cells is a promising avenue of research, these predictions must undergo intensive *in vitro* and *in vivo* tests to validate their therapeutic potential.

## CONCLUSIONS

In the past few years, a great deal of research has been directed towards uncovering the functional dependencies of (intra and inter) cellular communications using network inference. The availability of high-throughput data along with the advancement of web-based technologies has led to the development of network visualization tools to mine and explore large-scale interaction graphs. However, web-based applications to enable the visualization of bipartite gene co-expression networks are still lacking. To address this issue, we developed *CrosstalkNet*, a user-friendly web-application that can be used to explore and mine interactions between tumor epithelial and stromal cells. The application gives users the ability to upload their customized files and, to search and highlight a gene of interest in a dense network, along with zoom features and several layout options. Users can also explore their inferred bipartite networks by focusing on a specific subgraph of a network, filtering the network based on interaction strength, investigating the possible paths between two genes and identifying the genes with the highest number of interactions (hubs).

In the present study, we have described the use of *CrosstalkNet* with biologically relevant case studies reporting relevant epi-stroma interactions in ER+ breast cancer. The neighbor explorer panel was used to extract shared first neighbors between two interferon-induced proteins, IFIT1 and OAS1 sharing ~88% of their interactions enriched in interferon, immune signaling pathways are over-represented on these overlapping genes [24]. We found that the FGFR1 gene is involved in the paracrine interactions, which are a key contributor for tumor growth and an actionable target for breast cancer treatments [31]. We also showed that a key gene involved in breast cancer progression, S100A7, is gained in the tumor specific network [32]. Exploring the highly connected gene in the network we identified the BDNF gene as the largest hub (over 70 epi-stroma interactions), underlining its important role in tumor proliferation [33,34]. We also identified compounds that target these hubs in the network using CMAP. Overall, our case studies highlight the relevance and versatility of the *CrosstalkNet* web-application in cancer research.

The *CrosstalkNet* web-application can be improved in multiple ways. As future direction, we intend to incorporate the three following features: a) Community explorer - users will have the option to divide their bipartite networks into subnetworks (called communities) to enable higher-level visualizations of the network of epi-stroma interactions; b) Pathway analysis - enrichment of pathways on hubs of a network, or on a selected subnetwork/community using GO biological processes; c) Drug explorer - prediction of compounds that can target a given node in the network using *PharmacoGx* package [36].

In conclusion, *we developed CrosstalkNet*, a web-application enabling exploratory network analysis for researchers with no computational skills. *CrosstalkNet* will assist researchers across various disciplines along with the clinicians in mining complex networks to decipher novel biological processes in the tumor epithelial-stroma crosstalk, as well as in other studies of inter-compartmental interactions.

## METHODS

### Datasets

The transcriptional profiling data of tumor epithelial and stromal samples along with their normal counterparts were derived from published datasets. In order to obtain paired transcriptional profiling of tumor epithelial and stromal samples for each patient, laser capture microdissection was carried out on tissue samples, as depicted in Figure 7. Breast cancer datasets were processed and published in [4] (Supplementary Table 1). For this study, we focused on the ER+ breast cancer samples.

**Figure 7.**
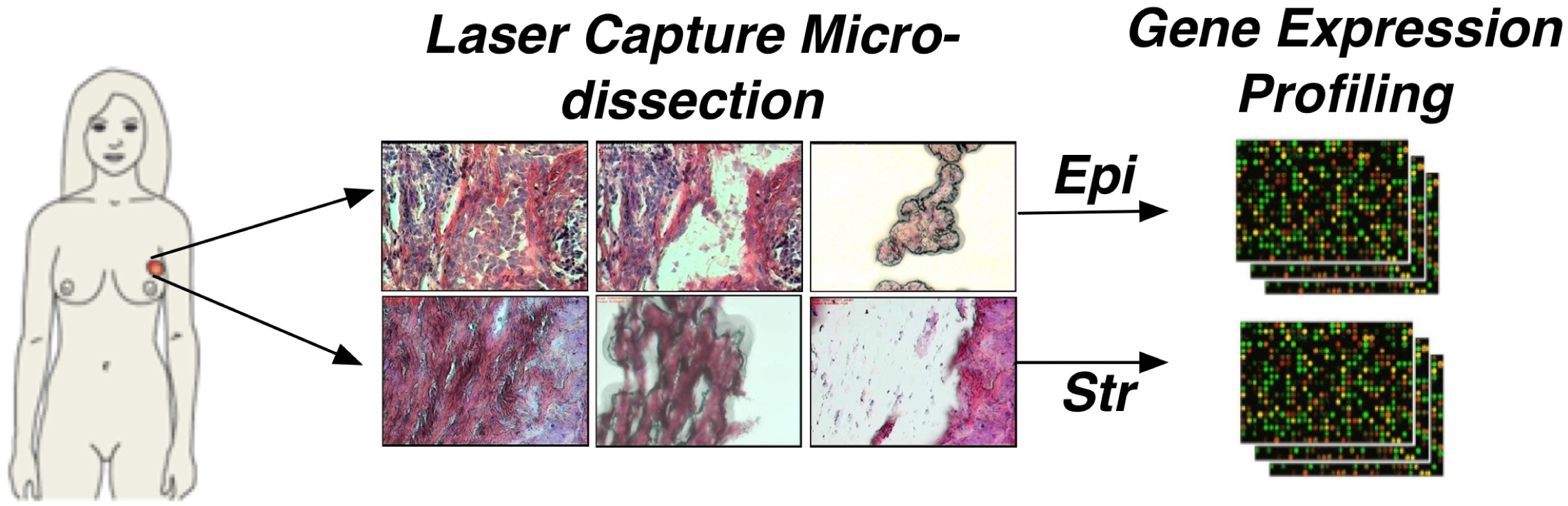
Gene expression profiling of tumor epithelial and stromal cells. The laser capturemicrodissection is carried out on tissue samples to obtain paired transcriptional ofiling of tumor epithelial and stromal samples for each patient.

### Network Inference

The genome-wide tumor epi-stroma network was constructed by estimating pairwise co-expression between the tumor epithelium and tumor stroma, while the normal network was constructed using the same approach based on normal epithelium and normal stroma. Gene co-expression was estimated using Pearson’s correlation coefficient. Supplementary Figure 1 depicts the computation of co-expression between *gene*_*i*_ in the epithelial cells with *gene*_*i*_, *gene*_*j*_, *gene*_*k*_ in the stromal cells. In the inferred network, each node represents the gene and each edge denotes a co-expression relationship between the epithelium and stroma. Depending on the type of stress, networks undergo condition-specific rewiring. The correlation based approach is extended by statistically comparing tumor microenvironment interactions inferred from tumor and matched normal samples. Such differential (delta) networks (defined as the difference between the tumor and normal epi-stroma networks) have proven to be efficient tools to investigate protein and genetic interactions, and are applied here for the first time to infer epi-stroma networks [10]. These differential networks give insight into the gain or loss of interactions in a tumor during carcinogenesis (Figure 2).

### Software Implementation

*CrosstalkNet* makes use of several technologies to achieve its results. The server uses node.js [38] and Rscript [39] in order to obtain graphs from the user’s query. Data are passed to and from R scripts via JavaScript Object Notation (JSON) in order to make the app have a minimal amount of dependencies. Different R scripts are used for different requests, and these scripts make use of a pool of data functions. Data models are defined in R to represent pseudo cytoscape.js [40] edges and nodes. Once the pseudo cytoscape.js edges and nodes are obtained from an R script, utility functions on the server add more properties to those objects and style them according to the layout specified in the request. This process leads to the creation of a final cytoscape.js configuration (config) complete with a layout, stylesheet, and elements. The config is sent to the client which will be responsible for rendering the graph. On the client side, cytoscape.js is used to display and interact with graphs. The client receives cytoscape.js configs from the server, adds event handlers for increased interactivity, and then displays the result to the user. The interface has several components separated into their own tabs. Structuring the interface in this manner reduces clutter while also making a clear distinction between the different tools and their responsibilities. The main technologies used for the interface are AngularJS and Angular Material [41]. Using AngularJS and Angular Material ensures a consistent look throughout the app in addition to making controls responsive. Furthermore, these technologies are well developed and maintained, making the app itself easier to maintain over time.

### User Access Levels

The data that is available to a user depends on the access level. Guest users have access only to sample data, while registered users have access to our processed data compendium in addition to their own uploaded files. Users are stored in a JSON file with their encrypted password. Once a user logs in, they are assigned a javascript web token (jwt) that will be then passed to the server on every request in order to identify that user.

### Performance

The performance of the app is a function of both the data file(s) being used as well as the query made to the server. This performance difference can be seen mostly in the main graph tab. The R script responsible for this can take up to 6-8 seconds to complete the execution since it may have to create more than 30,000 edges. Additionally, the client (i.e., the browser) also takes longer to load the graph due to the rendering of many elements. If more than 10,000 elements are shown in a graph, interacting with the graph becomes slow, since event handlers take longer to execute for a large amount of elements.

The interface can lag slightly when very large graphs are rendered, or when large tables are displayed. This is a drawback with AngularJS, since it has to keep track of changes on large collections of objects which is time consuming. One-time binding was employed in order to eliminate the more serious lag associated with large objects that remain static between requests made by the user. Other causes for interface lag can be attributed to Angular Material’s expensive animations and transitions.

### Features

The web-app has various features to mine and analyse large-scale interaction networks. Most of the important features of the application are described below.

- *Upload:* Users have the ability to upload their own adjacency matrix files to the server. Uploaded files must have an Rdata extension and must contain a single dgCMatrix (sparse matrix) from the *Matrix* package saved using the *saveRDS* function. Furthermore, row names and column names have to match, and the file must not contain missing (NA) values. Furthermore, row and column names and either be gene symbols or entrez gene id’s. Uploaded files will be private to the user that uploaded them and cannot be accessed by other users.
- *Main Graph:* The Main Graph is used to view the first and second neighbors of multiple genes of interest at the same time. Viewing the interactions this way makes it easy to see the neighbors that a certain set of genes have in common thereby revealing pathways between these genes in an intuitive way. To prevent graphs from getting unmanageably large in size, a filtering option is provided in order to allow filtering of both first and second neighbors individually. This makes it possible to focus on the more relevant interactions. There is a table view showing the interactions between the genes in the corresponding graph, and the table is downloadable as a CSV file.
- *Elements and Network Styles:* There is no layout that is the best for every graph. *CrosstalkNet* gives multiple layout options for the end user. Furthermore, the information that can be inferred from a graph is very limited unless there is some way to distinguish between the strengths of interactions. The dynamic edge-styling done in *CrosstalkNet* makes it possible to discern strong interactions from weak interactions via gradient colors and edge widths.
- *Interaction Explorer:* Exploring beyond the level of second neighbors potentially allows for the discovery of more complex pathways. The interaction explorer enables this functionality by providing the ability to view arbitrarily many levels of neighbors.
- *Path Explorer*: The path explorer makes it possible to obtain all paths with a maximum of one hop between 2 genes of interest. This is particularly useful when wanting to analyze the strengths of the connections between these genes.
- *Degree Explorer:* Viewing genes ordered by their degree simplifies the process of determining which genes are potentially interesting. The degree explorer does this by showing either the top genes based on their degree, or by showing genes having a degree greater or equal to a specified number. It is capable of showing an arbitrarily large list of genes, which can be downloaded in a simple comma-separated value (CSV) format.
- *Edge inspector/Edge Highlighter:* The best way to mine data with the app is to interact with the graphs. Clicking on an edge will bring up a tool called the Edge Inspector which displays information about the edge such as its weight and the genes which it connects. This is particularly useful for delta networks where edge styling can’t say anything about the normal and tumor interaction, rather only the difference between the two.
- *Gene Locator:* The gene locator is an essential tool when large graphs are being displayed. Instead of manually scanning through the graph, a user can type a gene in a textbox with autocomplete which will indicate to them whether or not the gene even exists in the graph. If it does, clicking on the autocomplete suggestion will pan the graph to that gene and highlight it so that it stands out among the other genes.

### Availability of data and material

The examples used in the web application, namely, normal, tumor and differential epi-stroma networks can be downloaded directly from the tool. These examples are provided to guest users as well as registered users. The methodology used to construct differential network as well as the web-based tool can be used to mine networks based on pan cancer transcriptomic profiles.

### Ethics approval

No ethics approval was required for this study.

## Acknowledgements

We thank Gary Bader and Max Franz for their help in using the cytoscape.js framework that has been used to develop the *CrosstalkNet* web-application.

## Supplementary Tables

**Supplementary Table 1:** Compendium of breast tumor and normal epi-stroma datasets

| GSE | Tumor Epi-Stroma samples | Normal Epi-Stroma samples | Platform |
| --- | --- | --- | --- |
| GSE 4823 | - | 22 | Agilent G4112A |
| GSE 5847 | 34 | - | Affymetrix U133A 2.0 |
| GSE 10797 | 28 | 5 | Affymetrix U133A 2.0 |
| GSE 14548 | 9 | 14 | Affymetrix U133X3P |
| GSE 35019 | 11 | - | Whole Genome DASL |

| Groups | Samples |
| --- | --- |
| ER+ | 54 |
| ER- | 28 |

## Supplementary Figures

**Supplementary Figure 1:**
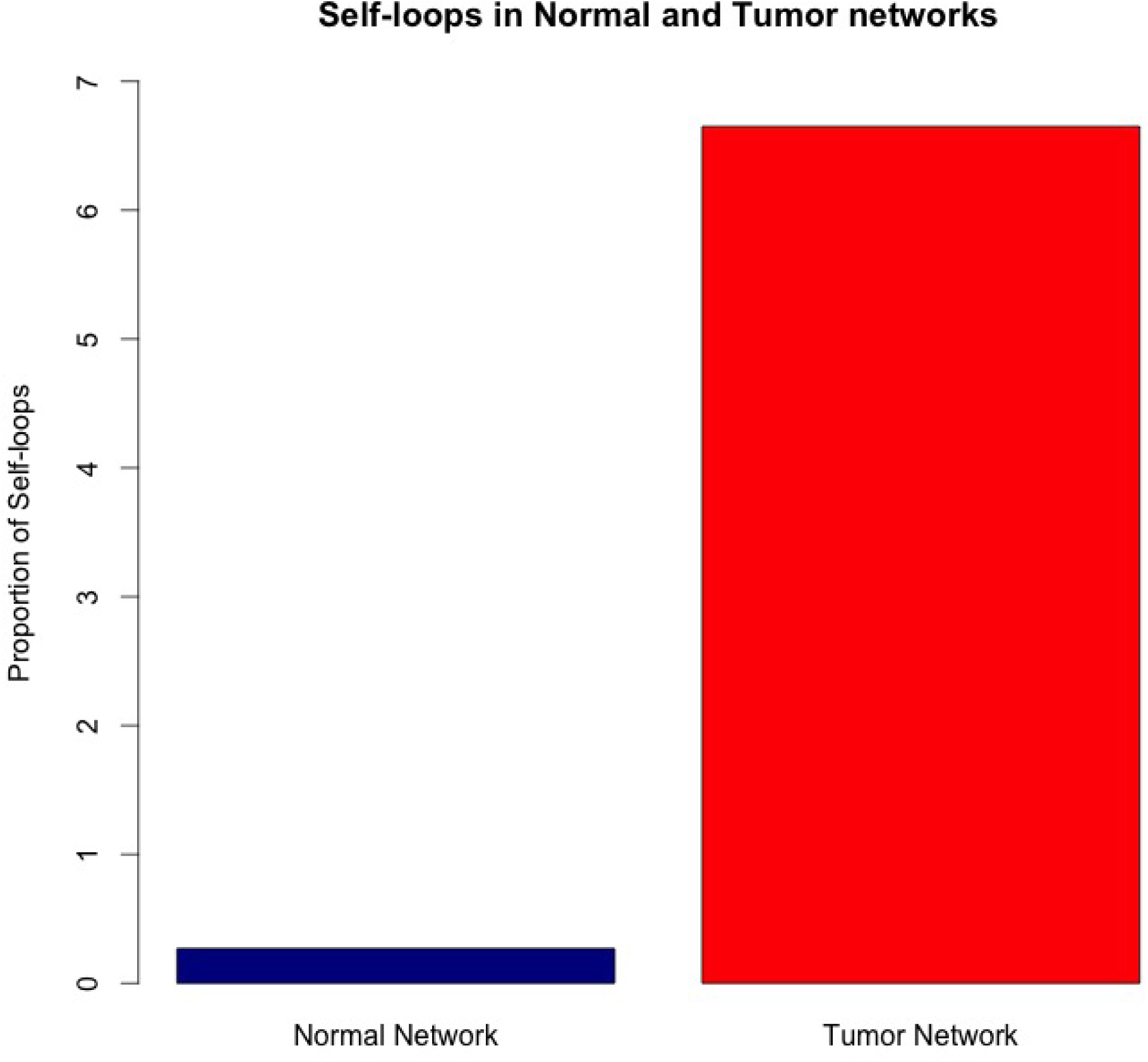
Proportion of self-loops in normal and tumor networks

**Supplementary Figure 2:**
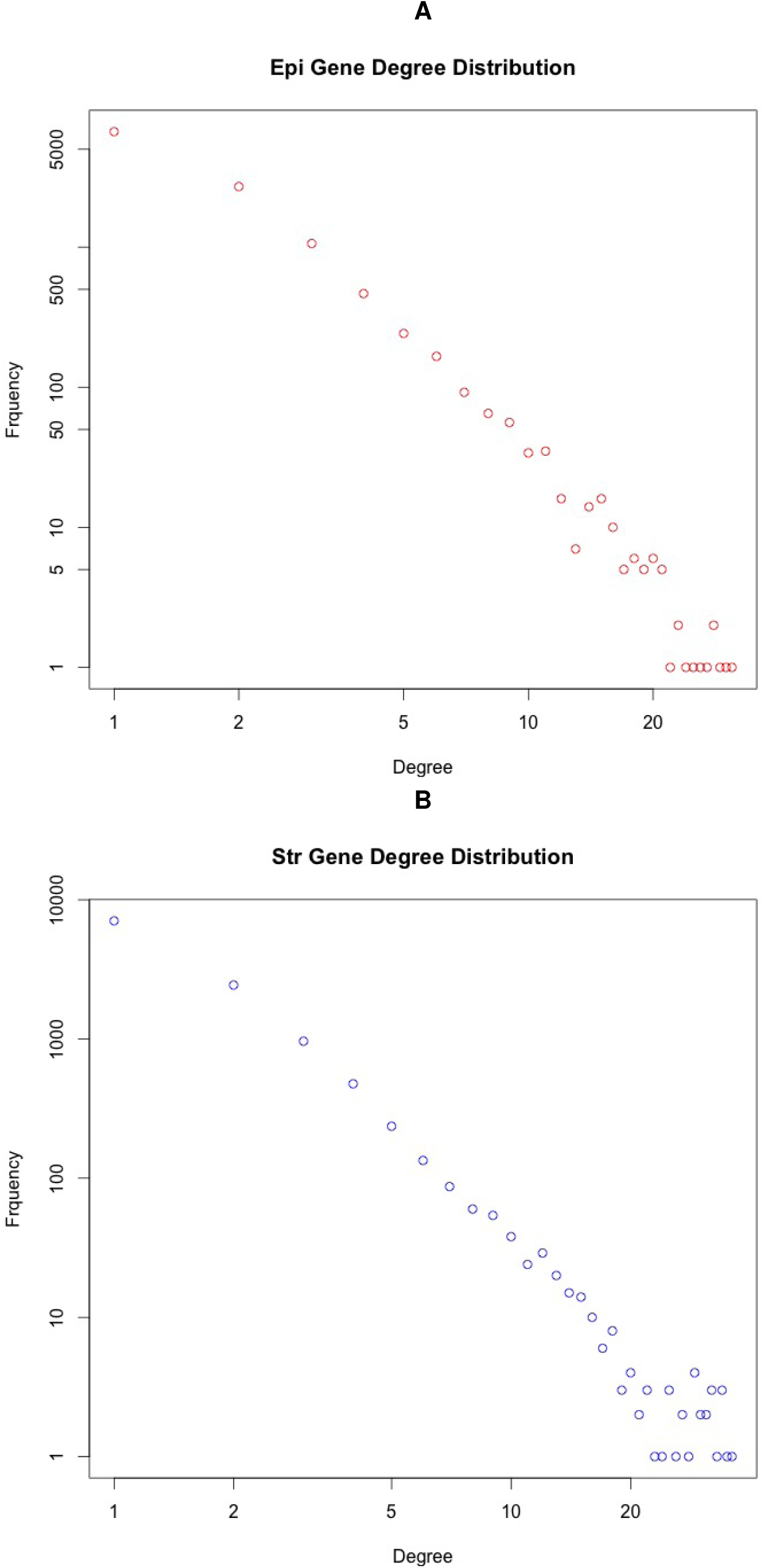
(A) Degree distribution of Epi genes in the tumor network (B) Degree distribution of Stroma genes in the tumor network

**Supplementary Figure 3:**
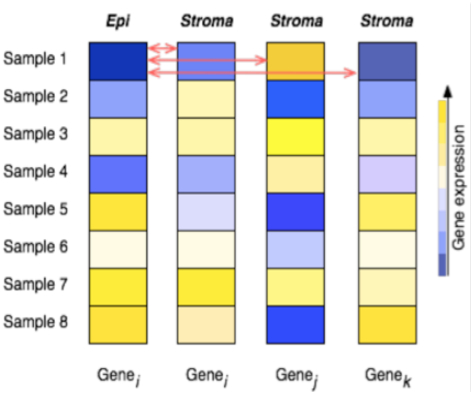
Construction of epi-stroma interaction network. A schematic diagram displaying the construction of the epi-stroma interaction network.

